# Examining historical rates of leafcutting bee cell pathogens to establish baseline infectivity rates for alfalfa seed growers

**DOI:** 10.1101/2022.01.09.475547

**Authors:** Justin Clements, Maggie Haylett, Brenda Nelson, Doug Walsh

## Abstract

The alfalfa leafcutting bee (*Megachile rotundata*) is one of the primary pollinators for the alfalfa seed industry. The alfalfa leafcutting bee is a solitary cavity nesting bee. Female *Megachile rotundata* bees will construct and provision individual brood cells lined with cut leaves (cocoon) and will gather nectar and pollen to place within the constructed cocoon. The female bee will lay a single egg within the constructed cocoon and leave the egg to undergo larval stage development and pupation into the adult stage. During this time multiple pathogens and parasitoids can prey on the developing larvae, resulting in the loss of the future adult bee. A major concern for commercial alfalfa seed growers is the presence of invertebrate pests and fugal pathogens. In the present study, we used historical data from the Parma Cocoon Diagnostic Laboratory to determine baseline rates of pathogen and parasite infection of *Megachile rotundata* cells and used this analysis to determine historical infection rates and cutoffs for management practices. Additionally, using a Faxitron (X-ray) analysis for *Megachile rotundata* cell obtained in 2020, we compared the presence of chalkbrood, pathogens, and parasitoids in samples collected from both growers’ stocks and newly purchased Canada bees. The results of the investigation demonstrate historical averages of the presence of chalkbrood, pathogens, and parasitoids. We also show a significant increase in chalkbrood and predators in 2007-2011 and a significant difference in chalkbrood and predators between bee samples obtained from Canada and grower stocks.

## Introduction

Alfalfa (*Medicago sativa*) is an important agricultural commodity with high production value throughout the United States [1,2]. Alfalfa, excluding other hay production, encompasses approximately 17 million acres of arable land within the United States, with a production value of over 8.8 billion dollars annually [1,3] and alfalfa production provides a vital resource for the livestock industry in the form of feed (hay and silage) [4]. Without the continued resource/value of this commodity, a stepwise lag in production of other commodities would be observed. For the continued success of alfalfa production, pollination and pollinator health is needed to sustain alfalfa seed crops [5]. Alfalfa seed production provides the germplasm to be used in the production of alfalfa hay. One of the primary seed producing regions in the world is the Pacific Northwest of the United States (Idaho, Oregon, and Washington) [6]. This agriculturally intense alfalfa seed production area is located in southwest Idaho, southeast Oregon (Treasure Valley), and southcentral Washington.

Alfalfa is self-incompatible and unable to self-pollinate, and an insect pollinator is necessary to generate a seed crop. As such, alfalfa seed producers rely heavily on pollination from the alfalfa leafcutting bee, *Megachile rotundata* F. (Megachildae) and the ground dwelling alkali bee (*Nomia melanderi*) [7–9]. *Megachile rotundata* bee is a solitary bee that does not form a beehive or produce honey. Instead, a single female *M. rotundata* will construct a nest that consists of individual brood cells that are lined with cut leaves (cocoon) [7,10]. Female bees will gather nectar and pollen and place the pollen within the constructed cocoon [7,11–12]. The female bee will lay a single egg within the constructed cocoon and seal the egg within the cocoon to undergo larval stage development and pupation into the adult stage [7]. The production of the cocoon and resources needed to provide nutrition to the developing larvae requires multiple trips to flowering plants, resulting in the highly desirable trait of a very effective pollinator species [2,7].

*Megachile rotundata* stocks can either be purchased from a commercial vendor or propagated by growers in bee boards within their own fields over multiple years [7]. One of the primary *M. rotundata* producing areas that growers purchase bee stocks from is in the central provinces of Canada, where *M. rotundata* are used to pollinate canola and alfalfa seed [7,13]. In these high latitudes with short growing seasons, *M. rotundata* will produce one brood per season. Commercially managed *M. rotundata* are provided with fabricated polystyrene foam bee boards to produce brood and nest cells. The foam bee boards are removed from the agricultural fields and transported to controlled cold rooms for storage. These nest cells are then sold to US alfalfa seed producers as first year stock [7]. It is generally considered that these bee stocks have a lower presence of invertebrate pests and fungal pathogens (e.g., chalkbrood) [7]. Supply and demand and currency exchange rates between the US and the Canadian dollar dictate the price paid for *M. rotundata* by US alfalfa seed growers. The health of *M. rotundata* bee broods are a large concern for US alfalfa seed growers since the purchase of *M. rotundata* can account for 20 to 40% of the operating expenses.

A major concern for commercial alfalfa seed growers is the presence of invertebrate pests and fungal pathogens within their bee stocks. During larval development within nest cells of *M. rotundata*, multiple pathogens and parasitoids can prey on the developing larvae, resulting in the loss of the future adult bee. Fungal pathogens include multiple *ascosphaera* species that result in the disease phenotype known as chalkbrood. Alfalfa seed growers are predominately concerned with the presence of *A. aggregate* within cells, as it is currently thought to be the predominate *ascosphaera* species which results in *M. rotundata* cell loss [14]. Besides fungal pathogens, nest cells are also predated on by multiple different parasitic wasp species including *Monodontomerus obscurus, Leucospis affinis, Pteromalus venustus*, and *Sapyga pumila* [15], nest destroying beetles including *Tribolium audax, Tribolium brevicornis, Trichodes ornatus* [15] and cuckoo bees (*Epeoloides pilosula*) [15]. The presence of these pathogens and parasitoids results in the loss of efficacy of growers’ bee stocks [2,15–16]. Additionally, these pathogens and predators can reproduce within grower stocks, bee boards, and housing, and if not controlled, can result in high abundance of dead bee larvae. Traditionally, growers monitor the presence of these pathogens using X-ray (Faxitron) imaging as a diagnostic technique [17]. In order to reduce cell loss, growers can use a combination of disinfectants and lures to protect bee cells from different pathogens and parasitoids. However, if stocks contain a high percent of any of these pathogens and parasitoids, growers are forced to burn/bury their current bee cell stocks, sterilize bee boards and housing, and purchase new bee cell stocks from commercial vendors [18–20]. Currently, the acceptable cutoffs for any of these pathogens are not well defined.

In the current investigation we examined archived data collected from the Parma Cocoon Diagnostic Laboratory from 1997-2021 to examine historical trends in the presence of pathogens and parasitoids infesting *M. rotundata* cells. These records provide baseline yearly infection rates of chalkbrood, predators, and parasites within historical samples and provide insights for growers regarding expectations and cutoffs for future *M. rotundata* stocks. We also examined the sex emergence ratio of *M. rotundata* cells to gain insight into the relative number of female bees emerging from cells, which are the primary pollinators of alfalfa seed [7]. Historical trends of pathogens and predators can provide insight into *M. rotundata* cell health and can provide valuable information on what can be considered as an appropriate baseline of infection for healthy bee stocks.

## Materials and Methods

### Data availability

All relevant data are contained within the paper and its supporting information files.

### Ethical statement

This article does not contain studies with any human participants and no specific permits were required for collection or experimental treatment of *Megachile rotundata* for the study described.

### Parma Cocoon Diagnostic Laboratory Archived Samples

The Parma Cocoon Diagnostic Laboratory is an extension orientated service that classifies the proportion of pathogen and parasitoid infected *M. rotundata* cells submitted by growers. Growers provide loose *M. rotundata* cells from individual populations to the diagnostic laboratory. From each population, five 10-gram samples are weighed and x-rayed encompassing approximately 500 bee cells per population. Each population is considered a single sample within this analysis. Archived records ranged from alfalfa seed production areas located in Idaho, Washington, Oregon, Montana, North Dakota and Canada. The service visually classifies fungal pathogens *Ascosphaera aggregata* and *Ascosphaera larvis*, insect parasites including imported chalcid wasps (*Monodontomerus obscurus*), cuckoo bees (*Epeoloides pilosula*), woodboring chalcid wasps (*Leucospis affinis*), long-tongued blister beetles (also known as sunflower beetles, *Nemognatha lutea*), Canadian chalcid wasps (*Pteromalus venustus*), and red-marked sapygids (*Sapyga pumila*), and predators/nest destroyers including American black flour beetles (*Tribolium audax*), giant flour beetles (*Tribolium brevicornis*), and checkered flower beetles (*Trichodes ornatus*) of cells using X-ray imaging (Image 1). Diagnostic records from 1997 to 2021 were compiled and statistically analyzed using analysis of variance (ANOVA) conducted in KaleidaGraph with a Tukey’s post hoc test to generate a correlation between the response variable (five-year intervals) and independent variables (predators, parasites, chalkbrood) using an α = 0.05 to examine temporal differences in *M. rotundata* cell health. In 2020, samples received from growers were designated as grower stock or recently purchased bees from Canada and statistically analyzed using a student t Test for unpaired data with unequal variance conducted in KaleidaGraph. Additionally, the sex ratio of leafcutting bee emergence was statistically analyzed using analysis of variance (ANOVA) conducted in KaleidaGraph with a Tukey’s post hoc test to generate a correlation between the response variable (year) and independent variables (male and female) using an α = 0.05. No distinction in geographic location was made in the analysis except for the data collected in 2020, which was used to examine differences in the presence of predators, parasites, and chalkbrood in newly purchased Canadian bees and grower stocks.

**Image 1:**
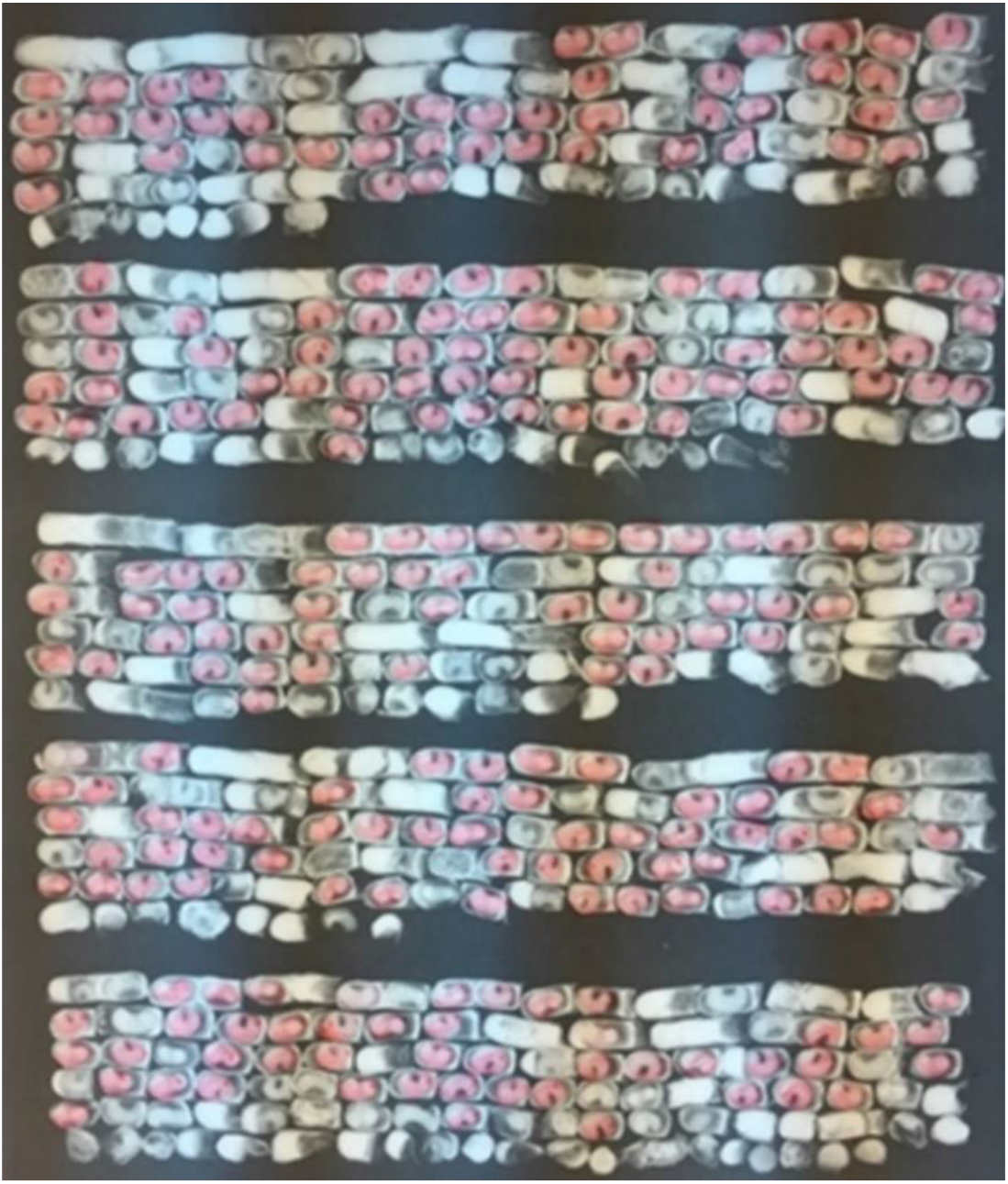
Faxitron X-ray of *M. rotundata* cells. Highlighted cells indicate healthy bee cells.

## Results

### Parma Cocoon Diagnostic Laboratory Archived Records

Compiled records from the Parma Cocoon Diagnostic Laboratory provide insight into the percent of chalkbrood, parasites, and predators found within *M. rotundata* cells processed at the diagnostic laboratory (**Figure 1**). The five-year average parasite infectivity rate and predator infectivity rate were not significantly different between most years examined and ranged from 3.10-4.99% and 0.72-7.821% respectively. From 1997-2001 there was a significantly higher infectivity rate of parasites compared to 2007-2011 (p=0.021). There was also significantly higher predator infectivity rate from 2007-2011 when compared to 1997-2001 (p < 0.0001), 2002-2005 (p < 0.0001), 2012-2016 (p= < 0.0001), and 2017-2021 (p= < 0.0001). The presence of chalkbrood was the most abundant in 2007-2011 when compared to 1997-2001 (p < 0.0001), 2002-2006 (p =0.0055), 2012-2016 (p= <0.0001), and 2017-2021 (p= < 0.0001) and five-year averages ranged from 3.27 – 10.65%. Within the 991 populations examined, the highest percent of chalkbrood found within an individual sample was 39% in 2012, the highest percent for a single sample of parasites was 40% in 2012, and the highest percent of predators was 28% in 2009. When comparing the yearly averages of chalkbrood, predators, and pathogens, infection with chalkbrood was statistically more abundant (p<0.0001) than both predators and parasites and, likewise, parasites were statistically (p < 0.0001) more abundant than predators when yearly averages were compared.

**Figure 1.**
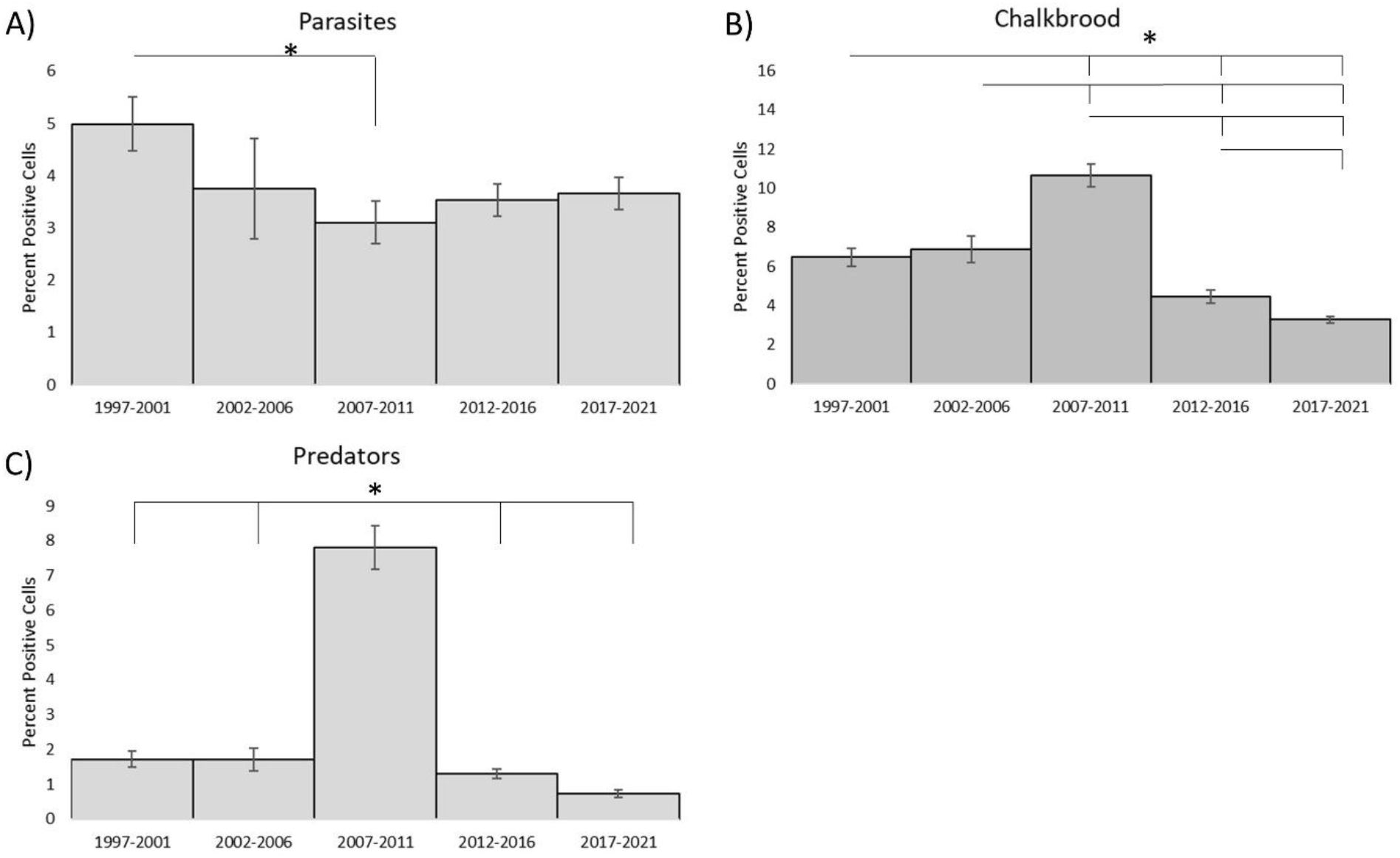
Parma Cocoon Diagnostic Laboratory *M. rotundata* archived records, A) percent parasites (*M. obscurus*, *E. pilosula*, *L. affinis*, *N. lutea*, *P. venustus*, and *S. pumila*), B) percent chalkbrood (*A. aggregate and A. larvis*), and C) percent predators (*T. audax, T. brevicornis, T. ornatus*). Significant differences are denoted with a line between treatments and an asterisk. Bars represent mean precent of infected cells within samples per year + standard error.

**Figure 2.**
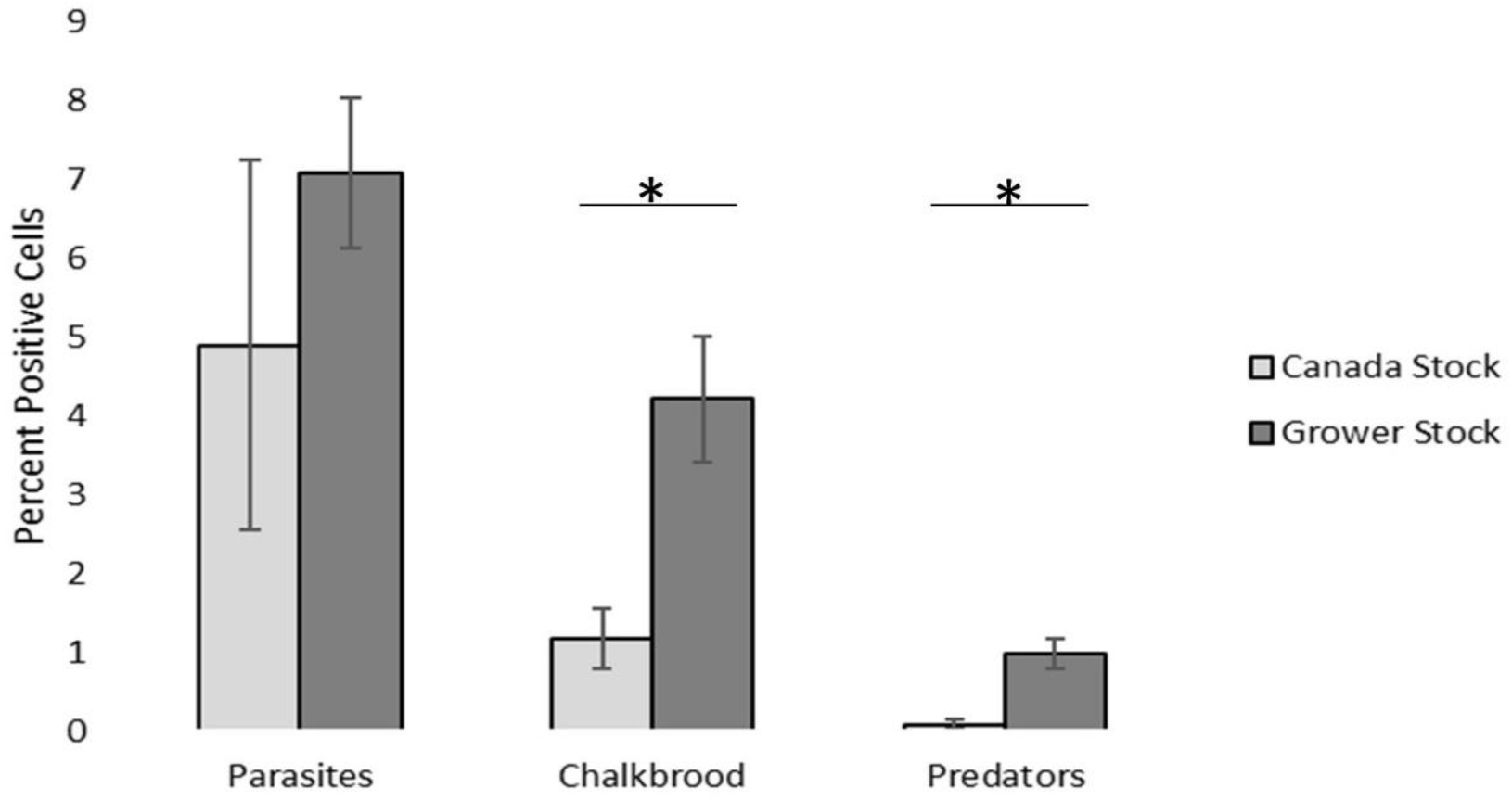
Difference between Canadian first year samples and grower stocks in 2020. Significant differences are denoted with a line between treatments and an asterisk. Bars represent mean precent of infected cells within samples per year + standard error.

### Grower Sample vs Canadian Bees

In 2020 we received 49 samples from growers to be examined for the presence of chalkbrood, parasites, and predators. From records provided by the growers, samples were designated as grower stock (n=43) or newly purchased bees from the central providence of Canada (n=6). When we examined newly purchased bee cells from Canada and grower stocks, we observed that the Canadian bees had significantly less chalkbrood (P=0.0011) and predators (P=0.00011).

### Sex Emergence Ratio

Archived records of the sex emergence of *M. rotundata* were also analyzed from 2010 to 2019 based on available data (**Figure 3**) to determine the ratio of female to male bees within the samples. No sex emergence ratios were available for 2013. Within all the years examined, there were statistically more male bees emerging than females with the exception of 2011 and 2012. Overall, when all samples between 2010-2019 were combined, there were statistically (P<0.0001) more males emerging in each sample (57.09%) compared to females (42.90%).

**Figure 3.**
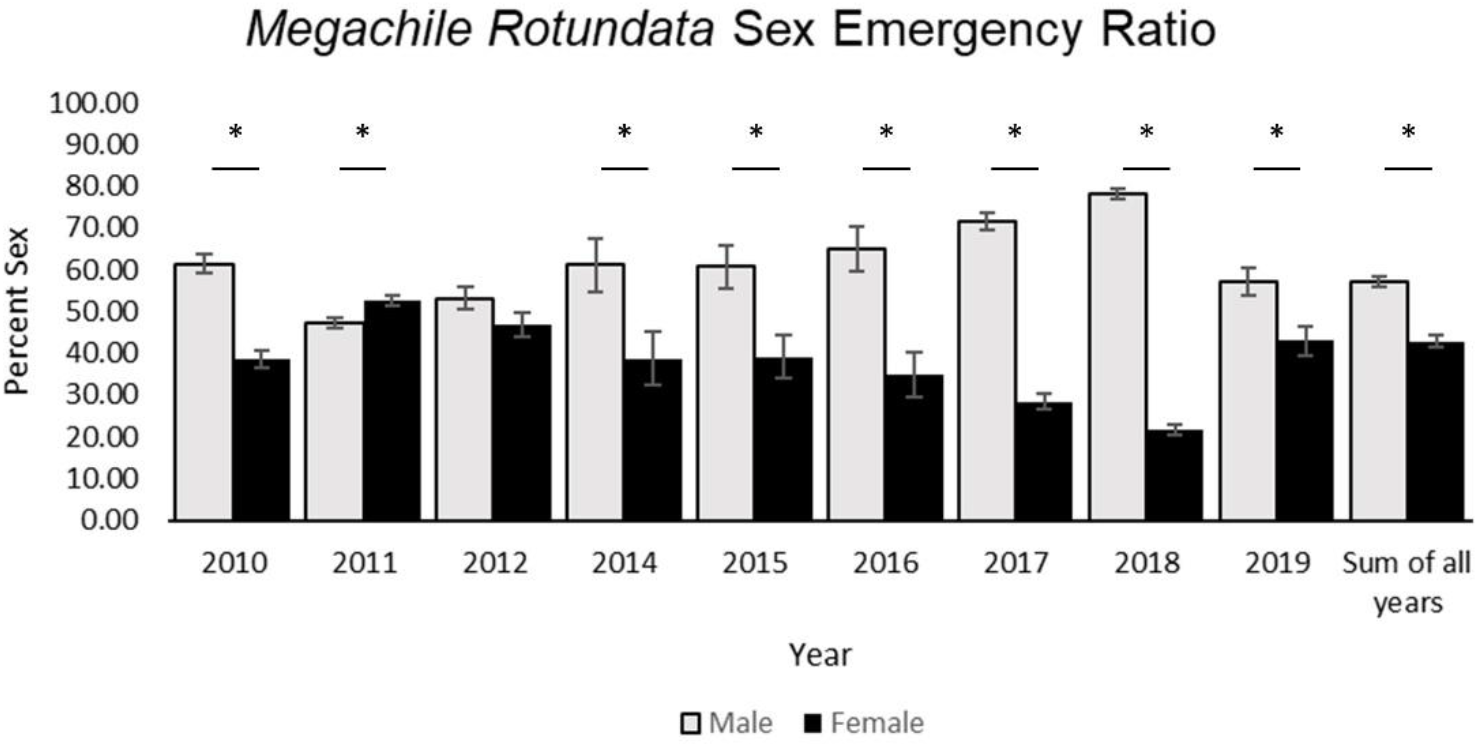
*Megachile rotundata* sex emergence ratio between 2010-2019. Yearly sex ratios that are significantly different are represented with asterisks. Bars represent mean percent male and female within a sample for each year + standard error.

## Discussion

The production of alfalfa seed is vital for the continued success of alfalfa hay and the livestock industry. Since alfalfa cannot self-pollinate, alfalfa seed producers rely predominately on two pollinator species, *M. rotundata* and *N. melanderi*. Unfortunately, *M. rotundata* cells play host to multiple different pathogen and parasitoids that can reduce the efficiency of grower bee stocks. Multiple investigations have explored and classified multiple different fungal pathogens, insect parasites, and nest destroying beetles [21–24]. While multiple different species that predate on *M. rotundata* cells have been determined, the presence of new and different species that infest cells needs to be constantly examined, allowing growers and integrated pest managers to develop new approaches to keep grower stocks healthy. This should include examining historical records to determine baseline percentages of pathogens, parasites, and predators which can be expected within samples. In the current investigation, we compiled archived records from the Parma Cocoon Diagnostic Laboratory to reveal trends in the presence of pathogens, parasites, and predators.

A principal goal of this investigation was to examine historical trends in the presence of pathogens, parasites, and predators from the Parma Cocoon Diagnostic Laboratory to understand historic normal ranges and to better inform growers and pest managers of expected and appropriate concentrations of these different classifications within their *M. rotundata* stocks. The diagnostic laboratory examined 991 historical samples encompassing ~590,000 cells to examine trends in *M. rotundata* health. Within the archived records, no mention of the specific pathogen, parasite, or predator species was made. Examining yearly averages, we noted that there was between 3.27 – 10.65 % of cells infected with chalkbrood, 0.72-7.81% infected with predators, and 3.10-3.75% infected with parasites. These mean averages can be used as a baseline for expected infection rates of samples and to inform growers regarding the health of their bee stocks. Besides the average infection rates that can provide insight for growers regarding expectations and cutoffs for future *M. rotundata* stocks, two important and significant observations can be drawn from this data. The first is that first year stocks of Canadian bees have a lower presence of chalkbrood and predators than grower raised bees that are propagated over multiple generations. While this data was only collected over one growing season, the findings support the regular purchase of new bee stocks to maintain bee health. The second is that there was statistically more chalkbrood and predators in bee cells from 2007-2011. Interestingly, the Canadian dollar increased in value from 2002-2007 and stayed on par with the US dollar though 2012 [25]. The exchange rate would have significantly affected the price of bees, resulting in higher grower cost which may have indirectly resulted in growers purchasing fewer bees from Canada. We hypothesize that growers would have propagated more bee stocks over this time period and not subsidized their stocks with newly purchased bees to cut input cost, which may have resulted in higher infection rate of both chalkbrood and predators in bee cells.

Within the current study, we determined expected presence of pathogens over all samples processed and found average infection rates of chalkbrood (5.54%), predators (2.32%), and parasite (3.74%) within historical samples collected at the Parma Cocoon Diagnostic Laboratory. While these values are only averages, they can provide insight for growers regarding expectations and cutoffs for future *M. rotundata* stocks. Knowing historical rates for percent of these pathogens and predators, growers can compare current bee stocks to historical samples. For example, the highest chalkbrood sample observed within this investigation was 39% infection rate, well above the 5.54% average and our extension recommendation would be to replace bees. Making these recommendations becomes more difficult when samples have infection rates closer to the average, but significant deviations are now easier to identify. In the current study, we did not investigate how infectivity rates affect pollination efficacy in the field. Future studies exploring pollination efficacy should be the focus of further investigation. As pollinators continue to be a vital resource for alfalfa seed producer, the agricultural community and growers should continue to monitor pollinator health and track trends in chalkbrood, predators, and parasitoids within *M. rotundata* stocks to make sure this important pollinating species remains a viable tool for alfalfa seed growers.

## Acknowledgements

This research was supported by funding from: United States Department of Agriculture Alfalfa Pollinator Research Initiative grant number 58-2080-0-009 awarded to JC. The authors would like to acknowledge Professor James Barbour for running the Parma Cocoon Laboratory before his retirement.

## Notes

### Competing Interest Statement

The authors have declared no competing interest.

